# Graph neural network–based prediction of direct reprogramming factors using gene regulatory networks with microRNA-mediated regulation

**DOI:** 10.64898/2026.01.28.702229

**Authors:** Ryota Kawasaki, Kazuhiro Takemoto, Momoko Hamano

## Abstract

Direct reprogramming (DR) converts somatic cells directly into target cell types while bypassing an intermediate pluripotent state, such as induced pluripotent stem cells. In practice, DR is achieved by transfecting multiple transcription factors (TFs); prior research has shown that combining microRNAs (miRNAs) with TFs further improves reprogramming efficiency. However, experimentally identifying effective TFs and miRNA combinations is difficult and costly, underscoring the need for robust *in silico* prediction approaches. We developed a graph neural network–based method to predict TFs that induce DR across diverse human cell types while explicitly modeling miRNA-mediated transcriptional regulation. By constructing a gene regulatory network integrating TF–target gene, TF–miRNA, miRNA–target gene, and gene–gene interactions, we implemented a Graph Attention Network v2 that predicts DR-inducing TFs while learning interaction importance and capturing transcriptional activation and repression. This approach outperformed existing methods in predicting experimentally validated DR-inducing TFs. Moreover, high-ranking predictions for previously unexplored tissues included TFs known to be associated with the development of the corresponding tissues, supporting the biological relevance of the results. Overall, the proposed method provides a practical in regenerative medicine.

## INTRODUCTION

Induced pluripotent stem cells (iPSCs) can differentiate into diverse cell types with specialized functions across the human body, avoiding the ethical issues associated with the use of human embryos inherent to embryonic stem cells ^1–3^. However, iPSC-mediated cell conversion requires weeks to months and carries a tumorigenic risk because cells pass through a pluripotent state ^2,4,5^. Direct reprogramming (DR) enables the direct conversion of somatic cells into target cell types while bypassing pluripotency ^6^. Compared with iPSC-mediated approaches, DR requires fewer gene transfections, thereby reducing tumorigenesis risk. Additionally, because somatic cells are not reprogrammed into iPSCs, their epigenetic state is largely preserved, further decreasing the time and cost required for cell generation ^7^.

DR-induced cell conversion requires transfection of specific combinations of key transcription factors (TFs). For example, mouse cardiomyocyte-like cells have been induced from postnatal cardiac or dermal fibroblasts using Gata4, Mef2c, and Tbx5 ^8^. However, experimentally identifying DR-inducing TFs across many cell types remains time-consuming and expensive. Consequently, several *in silico* methods have been developed to predict DR-inducing TFs. CellNet constructs gene regulatory networks (GRNs) and identifies TFs with high network influence ^9^. Entropy-based approaches predict DR-inducing TFs using the Jensen–Shannon divergence matrix ^10^. TRANSDIRE integrates multi-omics data while accounting for transcription with pioneer factors to predict candidate TFs ^11^, and ANANSE infers DR-inducing TFs by modeling enhancer-driven transcriptional regulation ^12^. Notably, these approaches focus primarily on TF-driven gene activation and do not consider gene regulation mediated by microRNAs (miRNAs).

As noncoding RNAs that typically bind complementarily to the 3’ untranslated region of target mRNA, miRNAs regulate gene expression through mRNA degradation or translational repression ^13,14^. Their importance in DR has been demonstrated in multiple studies ^15^. For example, combining Gata4, Mef2c, Tbx5, and miR-133a promotes DR from mouse or human fibroblasts to cardiomyocytes, as miR-133a directly represses Snai1, a master regulator of epithelial-to-mesenchymal transition ^16^. Similarly, cotransfection of miR-302 with Pdx1, Ngn3, and MafA promotes DR from hepatocytes to pancreatic cells by repressing the translation of multiple epigenetic regulators and inducing DNA demethylation ^17^. miR-9-5p/-3p and miR-124 induce neuronal conversion from fibroblasts by reconfiguring chromatin accessibility, promoting neuronal identify while suppressing fibroblast programs ^18^. Collectively, these findings indicate that miRNA-mediated posttranscriptional repression is critical for efficient cell conversion, underscoring the need to integrate TF-driven gene activation and miRNA-mediated gene regulation into predictive models of DR-inducing TFs.

In this study, we developed a novel computational framework to predict DR-inducing TFs across diverse human cell types while explicitly accounting for miRNA-mediated gene regulation. We constructed an integrated GRN incorporating TF–target gene, TF–miRNA, miRNA–target gene, and gene–gene interactions, enabling prediction of DR-inducing TFs by modeling gene activation and repression. The method achieved accurate and interpretable predictions for TFs that induce DR from fibroblasts into six cell types: cardiomyocytes, hepatocytes, pancreatic cells, neurons, Paneth cells, and skeletal muscle cells. To further demonstrate broad applicability, we generated predictions for 11 tissues lacking reported DR-inducing TFs and evaluated their biological plausibility.

## MATERIALS AND METHODS

### Gene Regulatory Interaction Data

Gene–gene interaction data were obtained from the Search Tool for the Retrieval of Interacting Genes (STRING) database (v12) ^19^ (https://string-db.org/) and the BioGRID database (v4.4.244) ^20^ (https://thebiogrid.org/). From STRING, which integrates physical protein–protein interactions and functional associations, we obtained 6,857,702 protein–protein interactions for humans. From BioGRID, which provides manually curated protein and genetic interactions in humans, we obtained 1,184,286 interactions.

From the miRTarBase database (v9.0) ^21^ (https://mirtarbase.cuhk.edu.cn/), we obtained miRNA–target gene regulatory data. This database provides experimentally validated human miRNA–target interactions, with 502,102 such interactions obtained in the present study. TF–miRNA regulatory interactions were obtained from TransmiR (v2.0) ^22^ (http://www.cuilab.cn/transmir), which curates TF–miRNA relationships from the literature and ChIP-seq data, yielding 588,626 interactions. Variants derived from the same precursor miRNA, such as miR-9-5p/-3p, were merged into a single miRNA entity.

TF–mRNA regulatory interactions were obtained from ChIP-Atlas 2.0 ^23^ (https://chip-atlas.org/), which contains >300,000 ChIP-seq, DNase-seq, ATAC-seq, and bisulfite-seq datasets. Combining the Target Genes function, which identifies genes regulated directly by specific TFs based on ChIP-seq binding profiles at specific genomic loci, with the hg38 reference genome, we identified TFs with MACS2 scores ≥50 binding within 5 kbp of target genes, resulting in 2,623,853 interactions.

### Gene Expression Profiles

Tissue-specific gene expression profiles were obtained from the GTEx Portal (v10) ^24^ (https://www.gtexportal.org/), which provides bulk RNA sequencing (RNA-seq) data from 54 tissues with 19,788 samples. We extracted transcripts per million (TPM)-normalized bulk RNA-seq data for 18 tissues, comprising 55,033 genes across 8,848 samples, including skin, heart, liver, pancreas, brain, intestine, skeletal muscle, kidneys, spleen, esophagus, lungs, thyroid, stomach, colon, adrenal gland, prostate, ovaries, and cervix samples.

### Integration of Gene Regulatory Interaction Data

Interaction data from all sources were first standardized to a two-column format labeled unified “Source genes” and “Target genes”. These datasets were then integrated to construct a unified GRN comprising 10,539,067 gene regulatory interactions (STRING: 6,857,702; BioGRID: 1,184,286; miRTarBase: 502,102; TransmiR: 588,626; ChIP-Atlas: 2,623,853).

The integrated GRN contained 24,523 unique genes, which were combined with 20,451 genes represented in the expression profiles to define a graph 𝐺 = (𝑉, 𝐸) with 24,763 nodes (genes), 𝑉 = {1,2, . . 𝑛 = 24,763}, and 10,539,067 edges (integrated interactions), 𝐸 ⊆ 𝑉 × 𝑉, where (𝑖, 𝑗) ∈ 𝐸 denotes an edge from node 𝑖 to node 𝑗. To uniformly handle heterogeneous interaction types, the network was converted to an undirected graph by symmetrizing the adjacency matrix 𝑨. If the adjacency matrix element 𝐴(𝑖, 𝑗) = 1, then both 𝐴(𝑖, 𝑗) and 𝐴(𝑗, 𝑖) = 1. The resulting graph had an average node degree of 429.76.

### Construction of Feature Vectors Based on Gene Expression Profiles

Tissue-specific expression profiles based on RNA-seq data were generated by averaging TPM values across samples within each tissue. From the initial 55,033 genes, we retained only those listed by the HUGO Gene Nomenclature Committee ^25^ as protein-coding genes (19,257 genes) or miRNAs (1,912 genes), yielding 20,451 genes.

Gene expression in target cells was represented by feature vectors 𝒙 = (𝑥_1_, 𝑥_2_, …, 𝑥*_n_*)^T^, and expression in the source-cell type (fibroblast) was represented by 𝒚 = (𝑦_1_, 𝑦_2_, …, 𝑦*_n_*)^T^, where 𝑥*_i_* and 𝑦*_i_* denote the expression levels of gene 𝑖 ∈ 𝑉 in the target and source cells, respectively. To quantify expression changes during reprograming, we calculated the log₂ fold change (log₂FC):

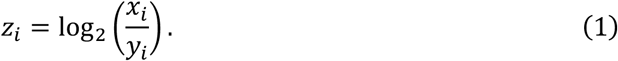

Values of 𝑧*_i_* were set to zero when 𝑥*_i_* = 0, 𝑦*_i_* = 0, or expression data were unavailable, thereby excluding extreme or biologically uninformative values arising from detection limits or missing data.

### Collection of Known DR-inducing Factors

Experimentally validated (known) DR-inducing factors were collected from published literature^7^ for six target cell types: cardiomyocytes (21 TFs), hepatocytes (18 TFs), pancreatic cells (10 TFs), neurons (58 TFs), Paneth cells (7 TFs), and skeletal muscle cells (6 TFs). The complete list of factors is provided in Supplementary Table 1.

### Construction of a Prediction Model Based on GATv2

Gene expression is governed by complex GRNs ^26^. TFs and miRNAs regulate gene expression in a cell-specific manner ^27,28^, and interactions among genes play central roles in DR ^29,30^.

We constructed a deep learning framework based on Graph Attention Networks v2 (GATv2) ^31^ to predict DR-inducing TFs. GATv2 is a graph neural network (GNN) capable of integrating gene expression profiles with GRN structure and addressing the static attention limitation of the Graph Attention Network (GAT) ^32^. By applying linear transformations and nonlinear activation functions after concatenating node features, GATv2 enables each node to dynamically weight it neighbors. We evaluated multiple GNNs and selected GATv2 because it achieved the highest prediction performance for DR-inducing TFs (Supplementary Figure 1) and provided interpretability by quantifying the importance of regulatory interactions in DR for each cell type through attention weights.

The model input consisted of the integrated GRN 𝐺(𝑉, 𝐸) and node-level gene features. We used log₂FC values 𝑧*_i_* (𝑖 ∈ 𝑉) as node features, thereby embedding cell type–specific gene expression changes into model training to predict DR-inducing TFs.

In the 𝑙-th GATv2 layer, 𝐾*_l_* attention heads were used. For each head 𝑘 (= 1, …, 𝐾*_l_*), an attention score 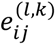 was computed between node 𝑖 and its neighbor nodes 𝑗 ∈ 𝒩*_i_*, quantifying the contribution of neighbor node 𝑗 to node 𝑖 based on their features 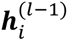 and 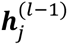. This score is expressed as follows:

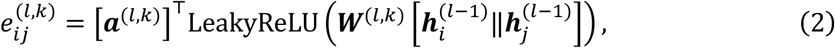

where 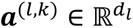 is the attention parameter vector (𝑑*_l_* is the feature dimension in the 𝑙-th layer), 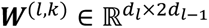 is the weight matrix for attention score calculation, and both are learnable parameters. The operator ‖ denotes concatenation.

Attention scores were normalized across all neighbors 𝑗 ∈ 𝒩*_i_*, using a softmax function to obtain attention coefficients:

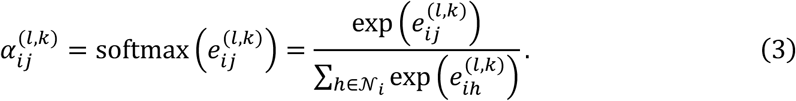

Using these coefficients, GATv2 computed the weighted averages of transformed neighbor node features:

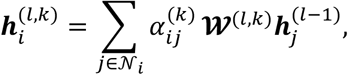

where 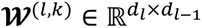 is a learnable weight matrix for feature transformation. Representations from all 𝐾*_l_* heads were concatenated and passed through an exponential linear unit (ELU) activation to produce the 𝑙-th layer output:

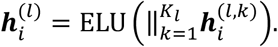

The proposed model architecture comprised two GATv2 layers followed by a fully connected layer. In the first layer (𝑙 = 1), we used 𝐾_1_ = 4 attention heads. In each head 𝑘, the feature 𝒉*_i_*^(0)^ = 𝑧*_i_* of each node was transformed into a 𝑑_1_ = 8-dimensional latent representation 𝒉*_i_*^(1,*k*)^. These 8-dimensional representations were concatenated to yield a 32 (= 𝐾_1_𝑑_1_)-dimensional intermediate representation. During training, dropout regularization was applied (rate = 0.2): 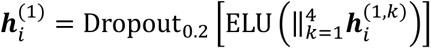.

In the second layer (𝑙 = 2), we used a 𝐾_2_ = 1 attention head (single-head) and transformed the 32-dimensional input into a 𝑑_2_ = 4-dimensional representation: 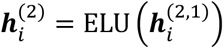.

In the final layer, the 4-dimensional node representation from the second layer was input into a fully connected layer, transformed into one-dimensional logits, and applied to a sigmoid function 𝜎(⋅) to calculate the probability that a gene induces DR, i.e., the DR induction probability: 𝑝*_i_* = 𝜎 (𝒘^T^𝒉*_i_*^(2)^ + 𝑏), where 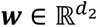 is the weight vector and b ∈ R is the bias term, with both being learnable.

The GATv2 model was implemented using PyTorch (v2.8.0) and PyTorch Geometric (v2.6.1). During training, binary cross-entropy loss was used as the loss function. To address class imbalance, the positive class weight (𝑝𝑜𝑠_𝑤𝑒𝑖𝑔ℎ𝑡) was set as the ratio of the negative example count 𝑁*_neg_* to the positive example count 𝑁*_pos_* (𝑝𝑜𝑠_𝑤𝑒𝑖𝑔ℎ𝑡 = 𝑁*_neg_*/𝑁*_pos_*). Model training used the Adam optimizer ^33^ with a learning rate of 0.01 for up to 100 epochs, with early stopping applied when the area under the receiver operating characteristic curve (AUC) failed to improve for five consecutive epochs. DR induction probabilities were calculated for 1,135 human TFs annotated in KEGG ^34^, and TFs were ranked in descending order of predicted probability.

### Prediction Performance Evaluation of the GATv2 Model via Cross-validation

Model performance was evaluated using leave-one-group-out cross-validation across six cell types with reported DR-inducing TFs: cardiomyocytes, hepatocytes, pancreatic cells, neurons, Paneth cells, and skeletal muscle cells. In each fold, one cell type served as the test set, whereas the remaining five types were used for training. For each test cell type, the corresponding log₂FC profile and the integrated GRN were provided as inputs. The integrated GRN was shared across all cell types to capture cell type–independent regulatory interactions. Using the log₂FC of each gene as node features corresponding to target cells, the model was trained to learn cell type–specific gene expression changes.

### Applicability of the Prediction Model to Tissues Without Reported DR-inducing TFs

To extend predictions to tissues lacking reported DR-inducing TFs, a final model was trained using all six annotated cell types. Predictions were then generated for 11 additional tissues: kidney, lung, thyroid, spleen, ovary, cervix, stomach, colon, esophagus, adrenal gland, and prostate tissues. For each tissue, log₂FC values relative to fibroblasts were computed and used as node features to calculate DR induction probabilities for each TF.

### Construction of TF–mRNA–miRNA Subnetworks Based on Attention Coefficients

Based on attention coefficients obtained during model training, subnetworks representing interactions around top-ranked predicted TFs were constructed to interpret and visualize prediction results. First, from each fold in the 6-fold cross-validation, attention coefficients 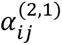 were extracted from the second GATv2 layer. The attention coefficients of all edges were sorted in descending order, and the top T = 100 highest-attention edge set 𝐸*_top_* was extracted. For each target cell 𝑐, the TF set 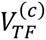 comprising the top 𝑁*_TF_* = 20 TFs by prediction score was identified, and neighboring genes connected by high-attention edges were extracted to construct a subnetwork 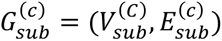, in which 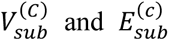 are expressed via the following equations:

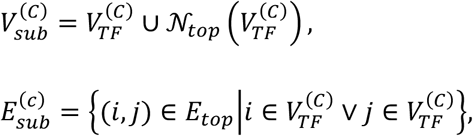

where 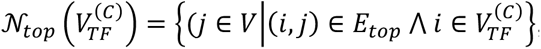, representing the neighbor node set of 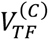 in 𝐸*_top_*.

### Visualization of TF–mRNA–miRNA Subnetworks

Extracted TF–mRNA–miRNA subnetworks were visualized using Cytoscape ^35^. Edge thickness reflected attention coefficients from the GATv2 model. To visualize gene expression level changes at each node during DR induction, log₂FC values relative to fibroblasts in each cell type were represented by node color, with red indicating upregulation (log₂FC > 0), green indicating downregulation (log₂FC < 0), and gray indicating unavailable expression data.

### Biological Function Evaluation of Top-ranked Predicted DR-inducing TFs

To assess the biological relevance of the top-ranked predicted DR-inducing TFs, Gene Ontology (GO) enrichment analysis was performed using the Database for Annotation, Visualization and Integrated Discovery ^36^ (https://davidbioinformatics.nih.gov/). Significantly enriched GO terms were identified using an adjusted p-value (*adjP*) of <0.05. For each cell type, enrichment results were evaluated to determine whether predicted TFs were associated with biological processes related to tissue development and differentiation.

## RESULTS

### Overview of the Proposed Method

We developed a GATv2-based machine learning framework that integrates gene expression profiles with a GRN, incorporating TF- and miRNA-mediated gene regulation, to predict DR-inducing TFs (Figure 1A–E). To quantify expression changes from source to target cells, we calculated log₂FC values using gene expression data, and these values were employed as node features in the graph (Figure 1A). We integrated gene regulatory interactions from five public databases to construct an integrated GRN comprising 10,539,067 interactions (Figure 1B). In addition, we manually curated known DR-inducing TFs from the literature for six cell types (cardiomyocytes, hepatocytes, pancreatic cells, neurons, Paneth cells, and skeletal muscle cells), which served as ground truth labels for model training and evaluation (Figure 1C).

**Figure 1.**
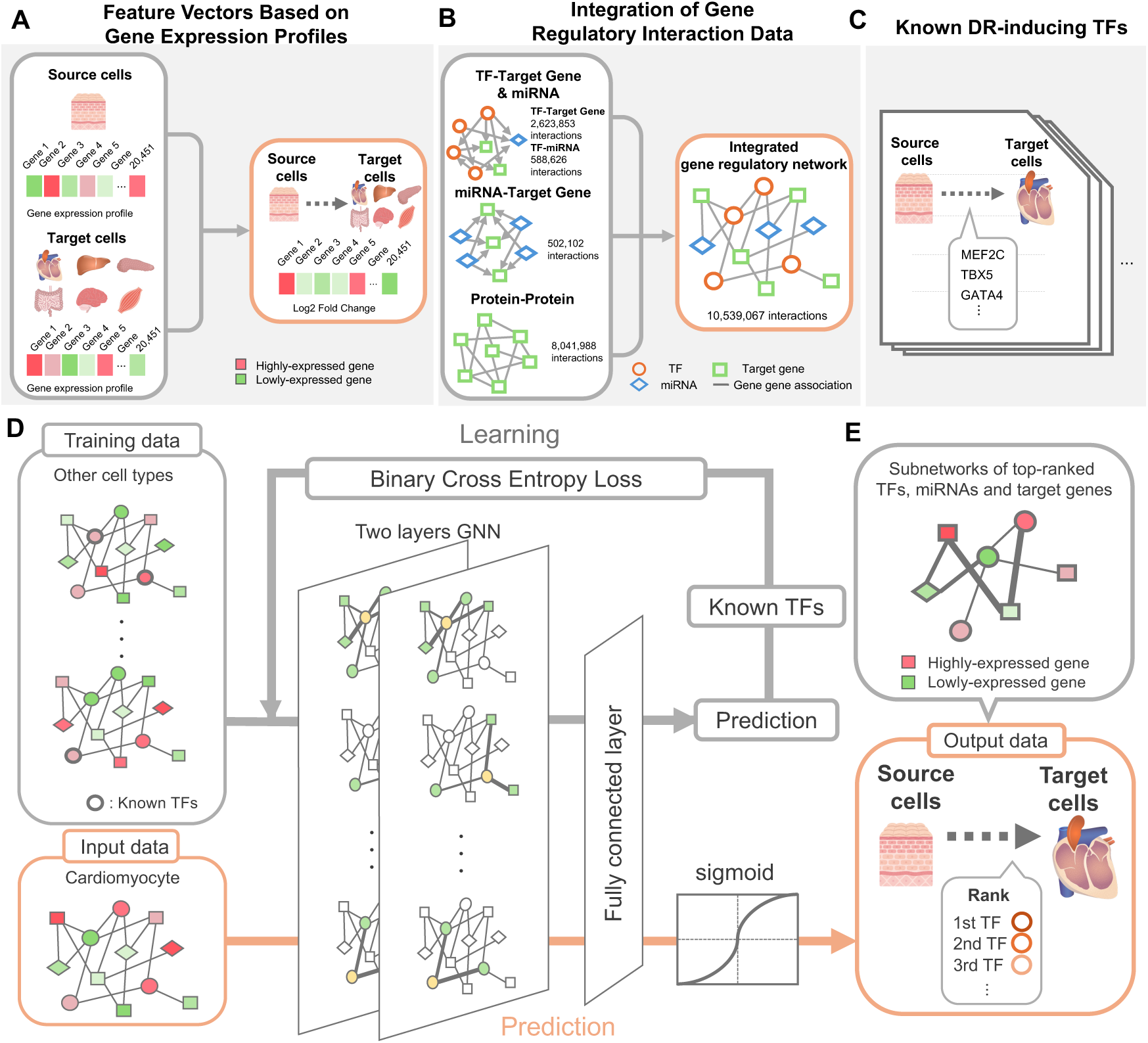
Input data and overview of the proposed method. (A) Gene expression profile data. RNA-seq data for fibroblasts, cardiomyocytes, hepatocytes, pancreatic cells, neurons, Paneth cells, and skeletal muscle cells were obtained from the GTEx Portal, and log₂FC values relative to fibroblasts were calculated for each target cell (20,451 genes). (B) Integrated GRN. Gene–gene interaction data were obtained from STRING and BioGRID, miRNA–target gene regulatory data from miRTarBase, TF–miRNA data from TransmiR, and TF–mRNA regulatory data from ChIP-Atlas. These datasets were integrated to construct a GRN containing 10,539,067 interactions. (C) List of TFs experimentally proven to induce DR, manually collected from the literature. (D) Overview of the DR-inducing TF prediction framework using a two-layer GATv2 with an attention mechanism. (E) TF–miRNA–mRNA subnetworks centered on top-ranked predicted TFs; attention coefficients were used to extract subnetworks that highlight important regulatory interactions among TFs, miRNAs, and target genes.

DR-inducing TFs were predicted using a two-layer GATv2 model. During training, the model assigned higher weights to important gene regulatory interactions through its attention mechanism and output a DR induction probability for each TF (Figure 1D). For the top-ranked TFs based on prediction score, we further constructed TF–mRNA–miRNA subnetworks using attention coefficients to visualize the key regulatory interactions underlying DR in each cell type (Figure 1E).

### Prediction Performance of the Proposed Method

We predicted candidate DR-inducing TFs for six cell types with known DR-inducing TFs and evaluated the model performance of each case (Figure 2A–F). The proposed approach, which integrates all interaction types (mRNA, TF, and miRNA), achieved a mean AUC of 0.865 across the six cell types (Figure 2A).

**Figure 2.**
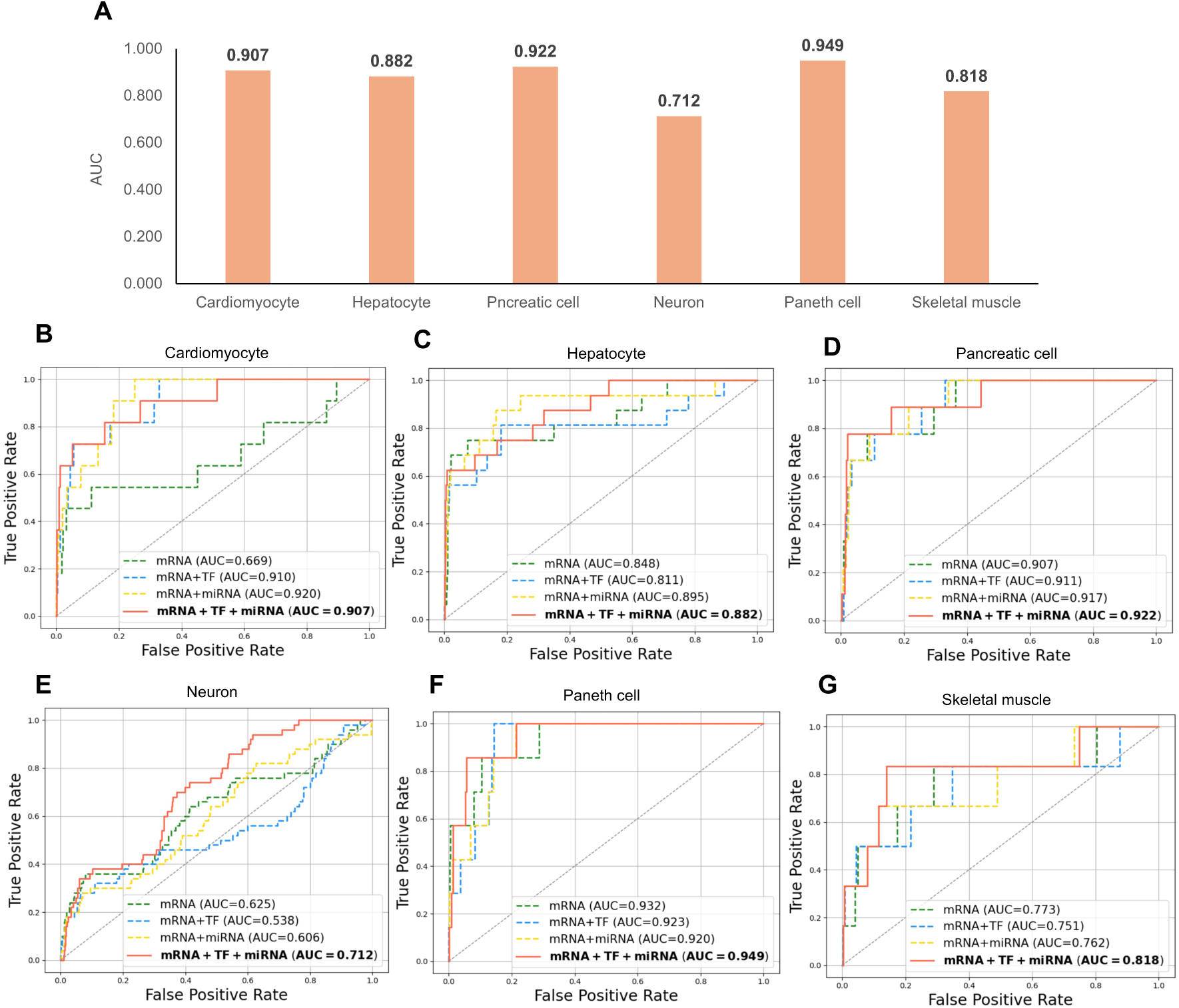
Prediction performance of the proposed method. (A) Prediction performance (AUC) of the proposed method applied to DR-inducing TF prediction across six cell types. (B–G) ROC curves showing differences in prediction accuracy when integrating multiple interaction types. ROC curves are shown for TFs predicted to induce DR from fibroblasts to (B) cardiomyocytes, (C) hepatocytes, (D) pancreatic cells, (E) neurons, (F) Paneth cells, and (G) skeletal muscle cells. Green dashed lines represent gene–gene interactions only, blue dashed lines represent gene–gene + TF–target gene interactions, yellow dashed lines represent gene–gene + miRNA–target gene interactions, and orange solid lines indicate gene–gene + TF–target gene + miRNA–target gene interactions.

Next, we evaluated the impact of different combinations of interaction datasets on prediction accuracy. Overall, prediction tended to improve as more diverse interaction types were incorporated across all cell types. Notably, inclusion of miRNA–target gene and TF–miRNA interaction data substantially enhanced performance. For example, in cardiomyocytes, the AUC increased markedly from 0.669 using mRNA interactions alone to 0.920 when miRNA interactions were added (Figure 2A). These findings indicate that explicitly modeling miRNA-mediated gene regulation improves the accuracy of DR-inducing TF prediction.

We also compared GATv2 with other GNN architectures (Supplementary Figure 1A–F), including graph convolutional network (GCN) ^37^ and GAT ^32^. GATv2 and GAT showed comparable performance in cardiomyocytes, hepatocytes, and Paneth cells (Supplementary Figure 1A, B, E), whereas in pancreatic and skeletal muscle cells, GATv2 achieved a markedly higher AUC compared with the next-best GAT model (Supplementary Figure 1C, F). When averaged across all six cell types, GATv2 achieved the highest AUC (0.865), followed by GAT (0.817) and GCN (0.724), demonstrating that GATv2 is a particularly effective architecture for predicting DR-inducing TFs.

### Prediction Results for DR-inducing TFs in Six Cell Types

We predicted DR-inducing TFs for reprogramming fibroblasts to six target cell types: cardiomyocytes, hepatocytes, pancreatic cells, neurons, Paneth cells, and skeletal muscle cells. Table 1 summarizes the top 20 TFs predicted to induce DR for each target cell type.

**Table 1.**
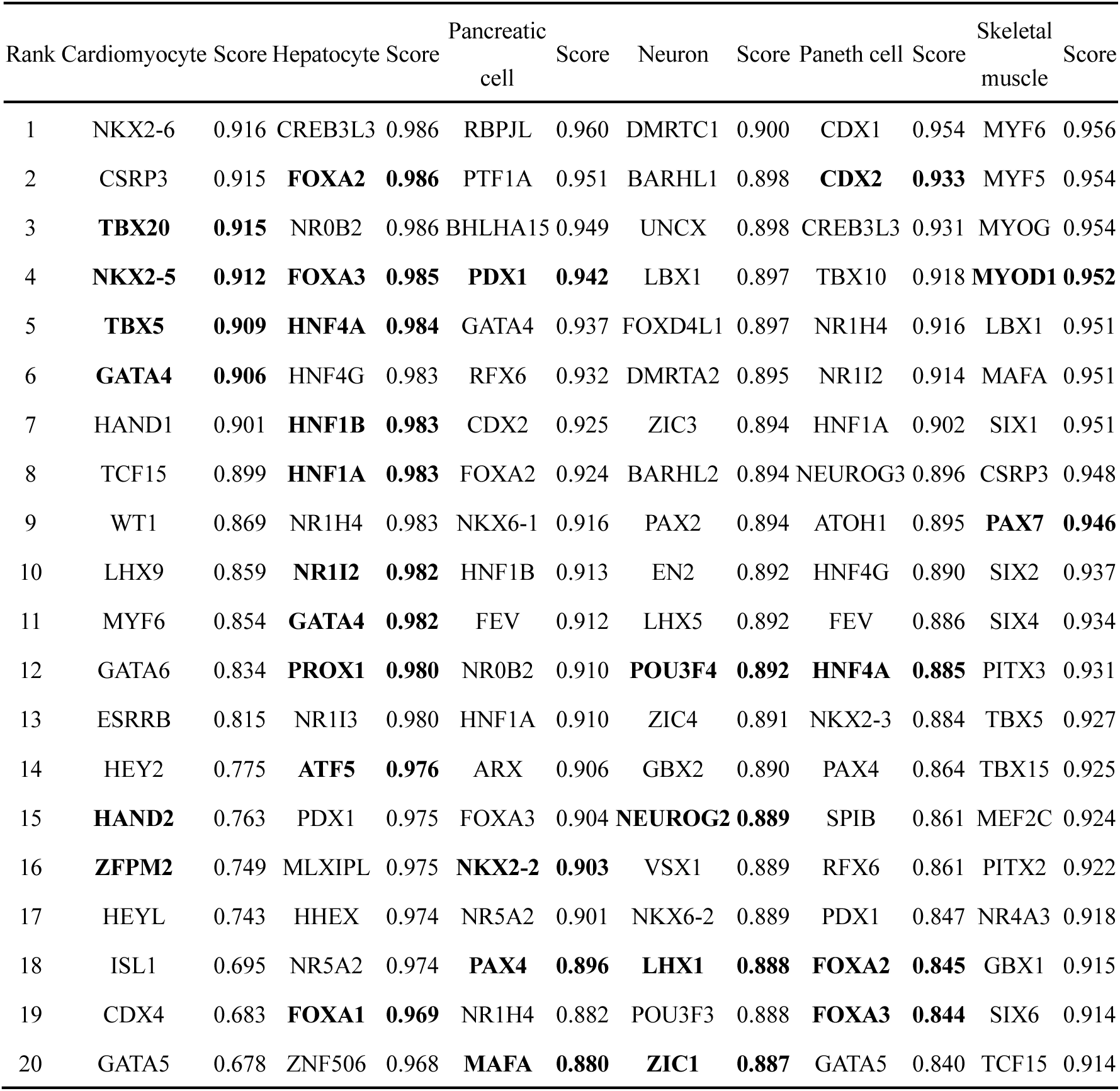
List of the top 20 TFs predicted by the proposed method for cardiomyocytes, hepatocytes, pancreatic cells, neurons, Paneth cells, and skeletal muscle cells (bold TFs indicate known DR-inducing TFs).

Next, we examined the overlap between predicted TFs and known DR-inducing TFs for each cell type. Among the top 20 predictions, the following known DR-inducing TFs were recorded: 6 for cardiomyocytes (38%), 10 for hepatocytes (56%), 4 for pancreatic cells (44%), 4 for neurons (8%), 4 for Paneth cells (57%), and 2 for skeletal muscle cells (33%).

Notably, among TFs not previously reported as DR-inducing, several highly ranked candidates directly regulate key genes in their respective cell types (Supplementary Figure 2). In hepatocytes, HNF4G binds the promoter regions of transthyretin ^38^, a liver-specific gene, and *AAT* ^38^, which is upregulated in human induced hepatocyte-like cells. In pancreatic cells, ChIP-seq data revealed that FOXA2 binds the promoter of *PAX6* ^39,40^, a gene required for pancreatic islet hormone gene expression during mouse pancreas development. Because these TFs directly regulate essential cell type–specific genes, they are likely involved in the molecular mechanisms underlying DR induction.

GO enrichment analysis of the top 20 predicted TFs revealed significant enrichment of biological functions related to the development and differentiation of each target cell (Supplementary Figure 3A–F and 4A–F). Together, these results demonstrate that the proposed method identifies biologically plausible DR-inducing TF candidates across diverse cell types.

### TF–mRNA–miRNA Relationships in DR for Six Cell Types

To evaluate the functional relevance of mRNAs and miRNAs interacting with top-ranked predicted TFs, we constructed regulatory subnetworks using attention coefficients derived from the GATv2 model. For each cell type, TF–mRNA–miRNA networks were visualized by extracting genes (TFs, mRNAs, and miRNAs) connected by the top 100 edges with the highest-attention coefficients, centered around the top 20 predicted TFs (Figure 3A–F).

**Figure 3.**
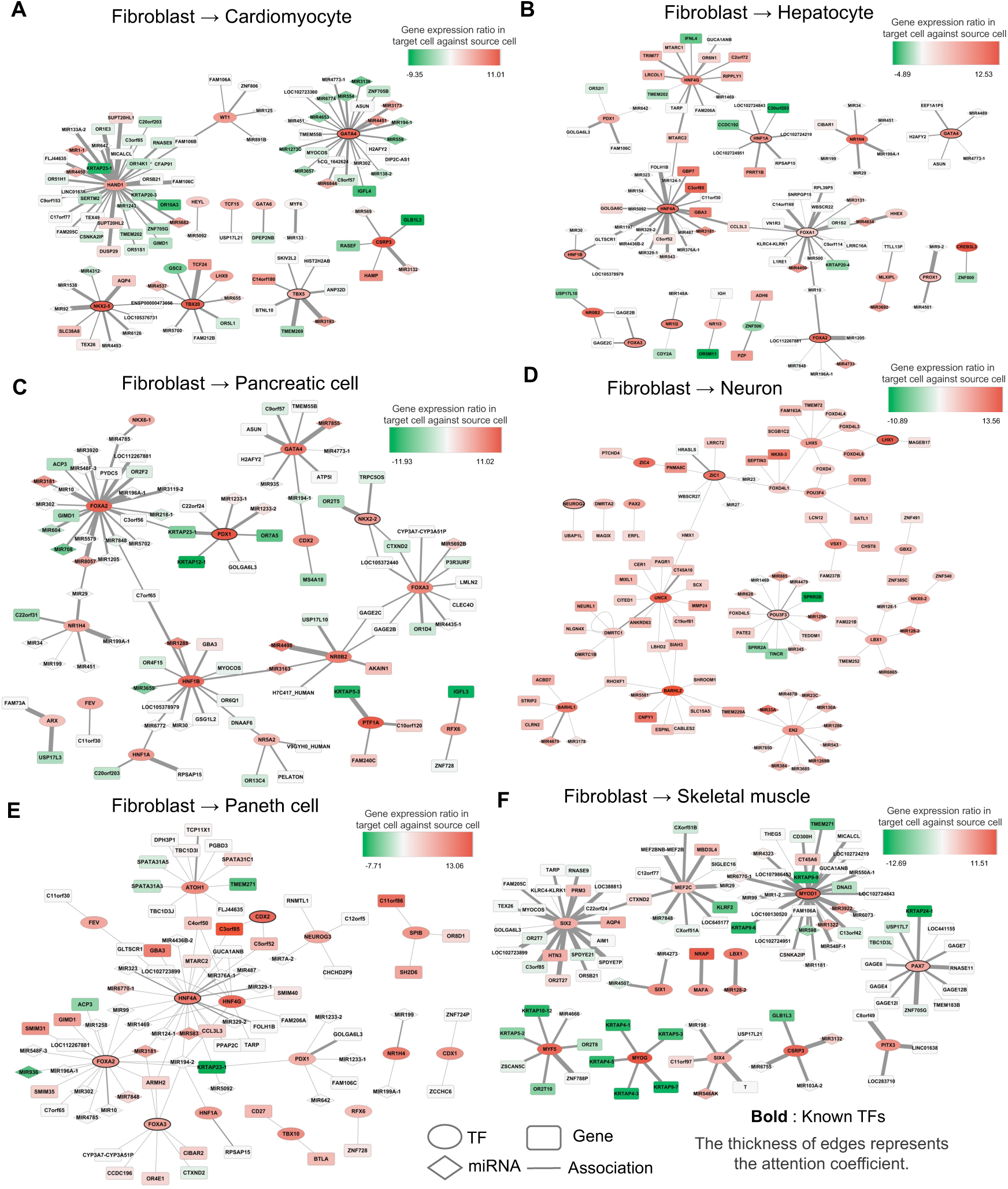
Importance of TF–mRNA–miRNA interactions predicted via the proposed method. GRNs of TFs predicted to induce DR from fibroblasts to (A) cardiomyocytes, (B) hepatocytes, (C) pancreatic cells, (D) neurons, (E) Paneth cells, and (F) skeletal muscle cells. Edge thickness represents attention coefficients derived from GATv2. Node color indicates gene expression changes (log₂FC) during DR induction from fibroblasts to each target cell type: red denotes upregulation (log₂FC > 0), green downregulation (log₂FC < 0), and gray unavailable expression data.

In cardiomyocytes, high-attention coefficients were assigned to interactions between HAND1 and miRNAs of the MIR133 and MIR1 families, which have been experimentally shown to promote DR ^16,41^, supporting the validity of our predictions. Notably, HAND1 exhibited more concentrated interactions relative to known DR-inducing TFs, such as GATA4, NKX2-5, TBX20, and TBX5. HAND1 is crucial for heart development, and its knockout causes defects in cardiac morphogenesis ^42^. Consistent with its association with heart development in GO enrichment analysis (Supplementary Figure 4A), HAND1 emerges as a promising candidate for cardiomyocyte DR.

In pancreatic cells, FOXA2 acted as a hub node comparable to known DR-inducing TFs (Figure 3C). FOXA2 regulates PDX1, a TF essential for initiating pancreatic development in the foregut endoderm ^43^, and its ablation results in hyperinsulinemic hypoglycemia, underscoring its importance for normal pancreatic β-cell function ^44^. Furthermore, interactions between FOXA2 and MIR302 were identified in the subnetworks; MIR302 has been experimentally shown to promote DR in pancreatic cells ^17^, suggesting cooperative regulation between TFs and miRNAs during DR. These findings support FOXA2 as a strong candidate for inducing DR in pancreatic cells.

Overall, these results demonstrate that newly predicted candidate TFs occupy central positions in GRNs and are functionally associated with tissue development. The attention mechanism of the GATv2 model enabled visualization of interactions reflecting cell type–specific regulatory mechanisms.

### Comparison of Prediction Performance with Previous Methods

Several computational approaches for predicting DR-inducing TFs have been reported previously. Therefore, we compared the prediction performance of the proposed method with CellNet ^9^, D’Alessio ^10^, TRANSDIRE ^11^, and ANANSE ^12^ (Figure 4A–F). ROC curves were generated for six cell types (cardiomyocytes, hepatocytes, pancreatic cells, neurons, Paneth cells, and skeletal muscle cells), and prediction performance was evaluated using the AUC. The proposed method achieved higher AUC values compared with the existing methods in cardiomyocytes, hepatocytes, pancreatic cells, and Paneth cells. In cardiomyocytes, the proposed method reached an AUC of 0.907, outperforming TRANSDIRE (0.839), ANANSE (0.835), and CellNet (0.724) (Figure 4A). In hepatocytes, it achieved an AUC of 0.882, outperforming TRANSDIRE (0.842), CellNet (0.784), and ANANSE (0.725) (Figure 4B). In pancreatic cells, the method showed the highest performance (0.922), with a marked improvement over ANANSE (0.625) and TRANSDIRE (0.760) (Figure 4C). In Paneth cells, the proposed method also achieved the highest AUC (0.949) (Figure 4E). For neurons and skeletal muscle cells, its performance was comparable to the best-performing existing methods (Figure 4D, F).

**Figure 4.**
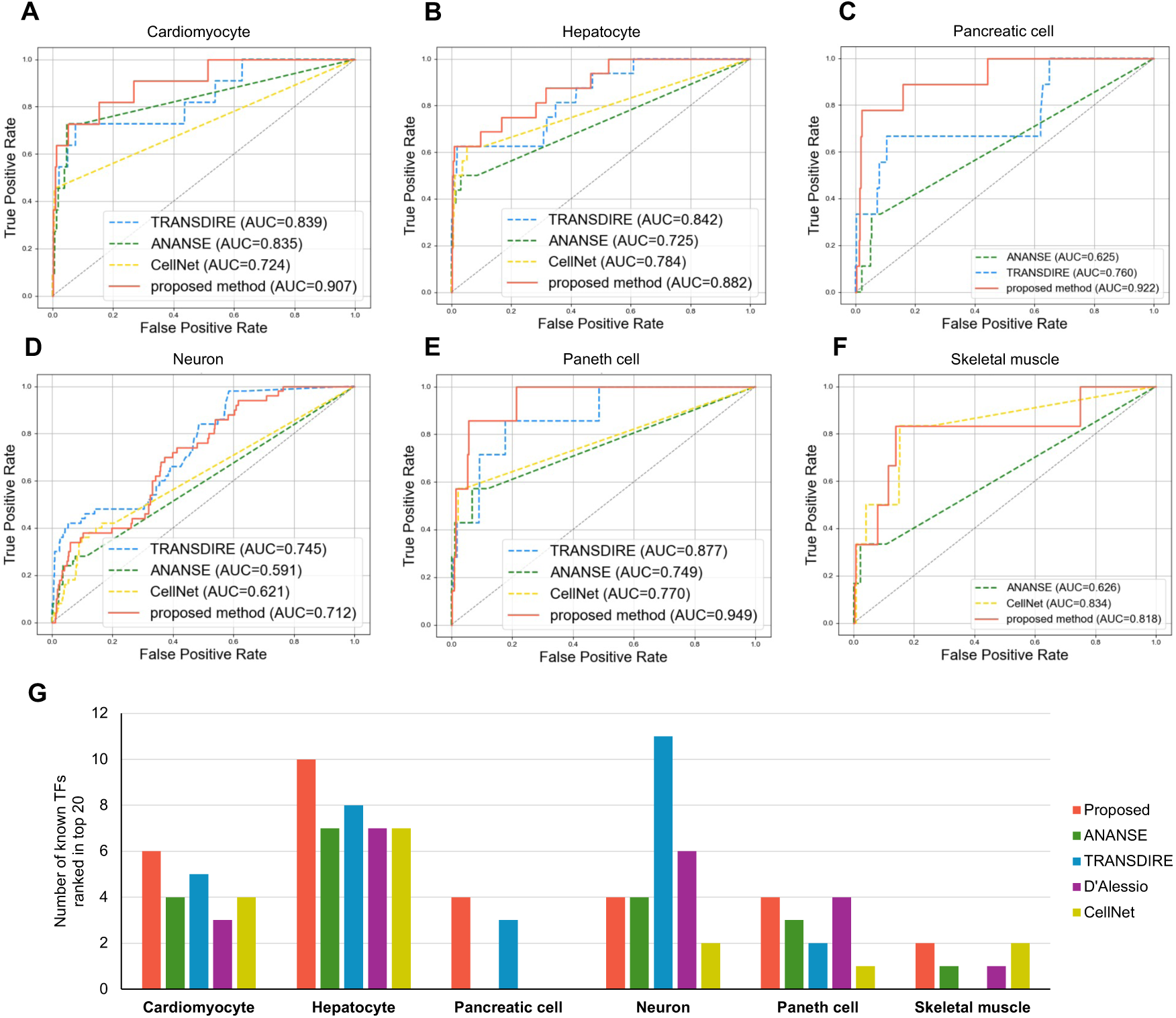
Proposed method’s prediction performance compared with existing methods. (A–F) Comparison of prediction performance between the proposed method and existing methods: TRANSDIRE, ANANSE, CellNet, and D’Alessio. ROC curves for TFs predicted to induce DR from fibroblasts to (A) cardiomyocytes, (B) hepatocytes, (C) pancreatic cells, (D) neurons, (E) Paneth cells, and (F) skeletal muscle cells. Blue, green, and yellow dashed lines indicate TRANSDIRE, ANANSE, and CellNet, respectively, whereas the orange solid line denotes the proposed method. (G) Comparison of the number of known DR-inducing TFs included in the top 20 predicted TFs. Blue, green, yellow, and purple bars indicate TRANSDIRE, ANANSE, CellNet, and D’Alessio, respectively, with the orange bar denoting the proposed method.

We further compared the number of known DR-inducing TFs among the top 20 predictions for each method (Figure 4G). The proposed method identified an equal or greater number of known DR-inducing TFs across all cell types except neurons. For example, in hepatocytes, the proposed method predicted 10 known DR-inducing TFs, outperforming ANANSE (7), TRANSDIRE (8), and CellNet (7). Overall, these results demonstrate that the proposed method provides superior or competitive performance relative to existing approaches.

### Applicability of the Proposed Method to Tissues Lacking Reported DR-inducing TFs

To assess the general applicability of the proposed method, we applied it to 11 tissues without reported DR-inducing TFs, i.e., kidney, lung, thyroid, spleen, ovary, cervix, stomach, colon, esophagus, adrenal gland, and prostate tissues. Using the proposed method trained on known DR-inducing TFs from six cell types (cardiomyocytes, hepatocytes, pancreatic cells, neurons, Paneth cells, and skeletal muscle cells), we predicted candidate DR-inducing TFs for these tissues (Figure 5).

**Figure 5.**
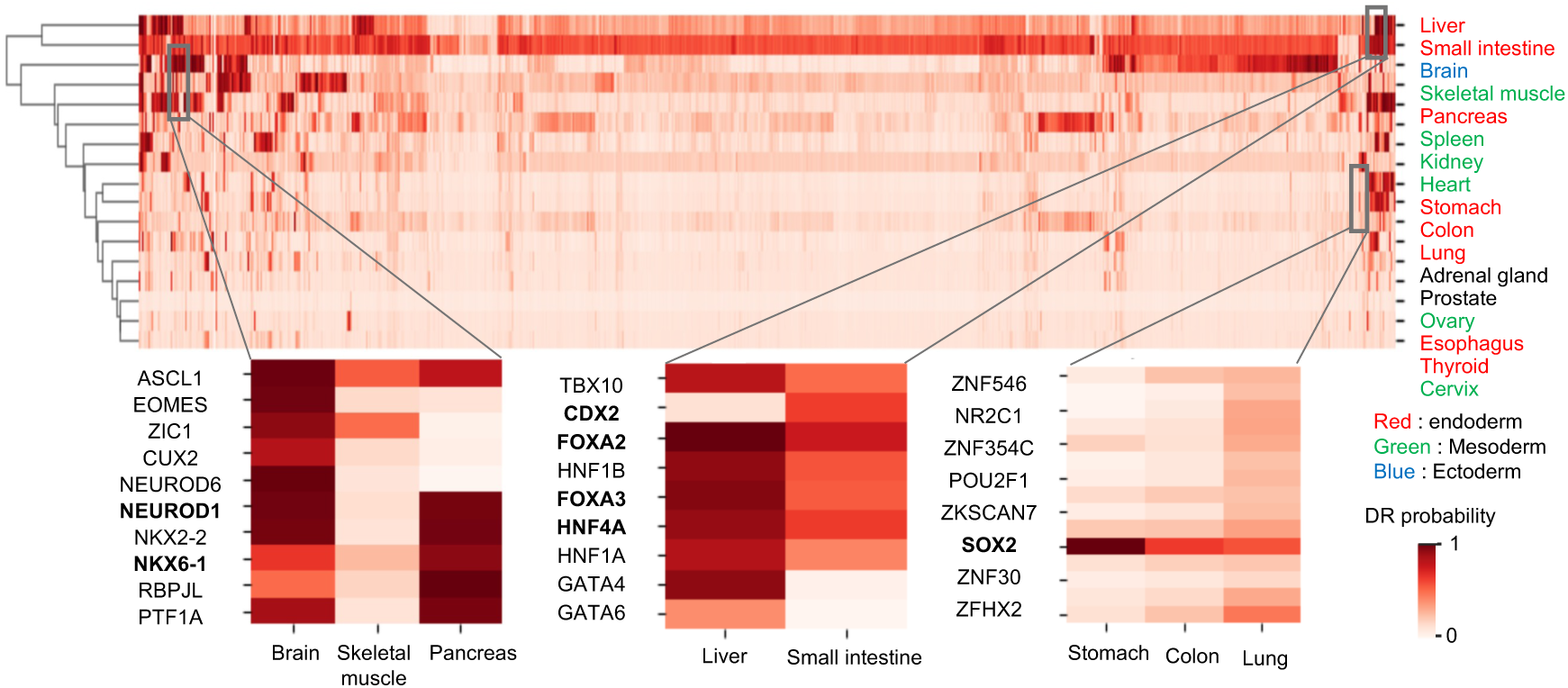
Applicability of the proposed method to tissues lacking reported DR-inducing TFs. Heatmap showing prediction scores for TFs across six tissues with reported DR-inducing TFs and 11 tissues without reported DR-inducing TFs (kidney, lung, thyroid, spleen, ovary, cervix, stomach, colon, esophagus, adrenal gland, and prostate tissues). Heatmap color indicates prediction scores calculated using the proposed method. Inset heatmaps highlight key TFs in related tissues. Cell types are color-coded by germ layer: red, endoderm; green, mesoderm; blue, ectoderm; tissues with multiple germ layers are shown in black.

In the colon, CDX1 and CDX2 ^45,46^ ranked first and second, respectively; these TFs are persistently expressed after the neonatal period in mice and are essential for anteroposterior patterning and intestinal epithelial development. In the thyroid, standard thyroid TFs (NKX2-1, PAX8, FOXE1, and HHEX) (56), commonly used as markers during thyroid differentiation induction from pluripotent stem cells, were ranked within the top 20 predictions (Supplementary Table 2). NKX2-1 is required to maintain normal thyroid architecture and function, and its overexpression induces the foregut endoderm toward a thyroid lineage ^47,48^. NEUROD1, a known DR-inducing TF shared between the pancreas and brain ^49,50^, received high prediction scores in both tissues. NKX6-1, which is selectively expressed in pancreatic β cells and subsets of neurons in adult mice ^51^, showed particularly high scores in the pancreas. FOXA family TFs and HNF4A, which are known DR-inducing TFs common to the liver and small intestine ^52,53^, were also highly ranked in both tissues. In contrast, CDX2, a known DR-inducing TF reported only in the small intestine ^53^, showed elevated scores only in this tissue. SOX2, which is required for gastric specification as it maintains chromatin accessibility at forestomach lineage loci ^54^, showed high scores in the stomach and lungs, both foregut-derived tissues, with particularly strong enrichment in the stomach.

To evaluate biological relevance, we performed GO enrichment analysis of the top-ranked TFs for each tissue. Results revealed significant enrichment for biological processes associated with tissue development (Supplementary Figure 5A–K). Collectively, these results indicate that the proposed method can identify TFs likely to regulate developmental programs even in tissues without previously reported DR-inducing factors.

## DISCUSSION

We developed a GATv2-based machine learning framework that predicts DR-inducing TFs by integrating gene expression profiles with a GRN. A novel aspect of this approach is the integration of miRNA-mediated gene regulation, enabling more comprehensive modeling of regulatory processes in DR and representing, to our knowledge, the first *in silico* method incorporating miRNA-mediated regulation for this task. Additionally, the proposed method offers high interpretability, as the GATv2 attention mechanism enables quantitative evaluation of regulatory interaction importance and visualization of TF–miRNA–mRNA subnetworks involved in DR. The method also demonstrated broad applicability by successfully predicting candidate DR-inducing TFs for tissues lacking previously reported factors.

Most existing DR-inducing TF prediction methods emphasize genes upregulated in target cells and the TFs that activate them. For example, CellNet focuses on TF-driven activation of highly expressed target genes. However, repression of specific genes is equally important during cell differentiation and reprogramming. Elevated miRNA levels regulate gene expression through mRNA degradation or translational repression and promote DR induction by repressing genes that maintain source-cell identity during lineage conversion ^16^. Accordingly, we incorporated miRNA–target gene and TF–miRNA interactions into the GRN. Furthermore, compared with single-cell RNA-seq, bulk RNA-seq can detect miRNA expression with higher sensitivity (Supplementary Figure 6A, B); therefore, bulk RNA-seq data were employed. Using log₂FC values derived from bulk RNA-seq as node features, the model learned regulatory patterns reflecting increased and decreased gene expression, enabling more comprehensive predictions of DR-inducing TFs that capture activation and repression mechanisms.

Another advantage of the proposed method is the integration of miRNA–target gene and TF–miRNA regulatory data, which allows prediction of miRNAs as DR-inducing factors, a feature not possible with previous approaches (Supplementary Table 3). For example, MIR384, associated with synaptic plasticity and brain development, ranked highly in neurons ^55^. In Paneth cells, MIR181A1, which regulates Wnt signaling and increases intestinal epithelial cell proliferation, was predicted ^56^. In skeletal muscle cells, MIR133B, a muscle-specific miRNA essential for proliferation and differentiation *in vitro* and *in vivo*, was highly ranked ^57^. These findings indicate that the proposed method can identify miRNAs critical for development and functional maintenance across cell types.

The proposed method enabled the identification of biologically plausible DR-inducing TF candidates for each cell type. Many top-ranked TFs have established roles in the development and function of these cell types. In cardiomyocytes, known DR-inducing TFs (such as NKX2-5, TBX20, and TBX5) appeared among the top five predictions. Attention-based subnetworks further highlighted HAND1 as a central hub with strong interactions with MIR133, a miRNA known to promote cardiomyocyte DR, suggesting HAND1 as a promising novel candidate. In hepatocytes and pancreatic cells, ChIP-seq data showed that HNF4G and FOXA2 bind promoters of cell type–specific and functionally important genes ^38–40^ (Supplementary Figure 2), implying roles in establishing target cell identity during DR. For tissues lacking reported DR-inducing TFs, developmentally critical TFs, such as CDX family members in the colon and NKX2-1 in the thyroid, were ranked highly, and GO enrichment revealed significant associations with tissue-specific developmental processes ^45–48^. Together, these results support the broad applicability of the proposed method and the biological relevance of its predictions.

In conclusion, the GATv2-based model accurately predicts DR-inducing TFs by integrating miRNA-mediated regulation with conventional TF–target gene interactions, advancing previous approaches by capturing both transcriptional activation and repression mechanisms critical for cell fate conversion. Nonetheless, this study has several limitations. Epigenomic features, such as DNA methylation patterns that influence cell fate transitions ^58^, were not incorporated. Additionally, metabolomic data reflecting cellular reprogramming dynamics were not included in our framework ^6,59,60^. Future analyses of multi-omics datasets with minimized batch effects across diverse cell types should enable construction of more comprehensive and accurate prediction models.

## ACKNOWLEDGEMENTS

The authors would like to thank Enago for the English language review.

## AUTHOR CONTRIBUTIONS

R.K. performed the preprocessing for public single-cell datasets, analyzed data and prepared the manuscript. M.H., and K.T. supervised the study, and R.K., M.H., and K.T. wrote the manuscript. All authors discussed the results and commented on the paper at all stages.

## FUNDING

This work was supported by a grant from JSPS KAKENHI Grant Numbers 22K18173 and 25K21559.

## Notes

### Competing Interest Statement

The authors have declared no competing interest.

