## Supplementary material for "Graph neural network–based prediction of direct reprogramming factors using gene regulatory networks with microRNA-mediated regulation": Supp. Table 1, Supp. Figure 1, Supp. Figure 2, Supp. Table 2, Supp. Figure 3, Supp. Figure 4, Supp. Figure 5, Supp. Figure 6, Supp. Table 3

### SUPPLEMENTARY DATA

**Supplementary Table 1.** List of TFs and miRNAs experimentally proven to induce DR.

| Target cell identity | Reported TFs or miRNAs for DR | PubMed ID |
|---|---|---|
| Cardiomyocyte | MIR208A | 22539765 |
| Cardiomyocyte | MIR499 | 22539765 |
| Cardiomyocyte | ETS2 | 22826236 |
| Cardiomyocyte | MESP1 | 22826236 |
| Cardiomyocyte | MIR1 | 23487791 |
| Cardiomyocyte | SRF | 23704920 |
| Cardiomyocyte | SMARCD3 | 23704920 |
| Cardiomyocyte | MEF2C | 23861494 |
| Cardiomyocyte | ESRRG | 24319660 |
| Cardiomyocyte | ZFPM2 | 24319660 |
| Cardiomyocyte | MYOCD | 24920580 |
| Cardiomyocyte | MIR133 | 24920580 |
| Cardiomyocyte | NKX2-5 | 26603904 |
| Cardiomyocyte | TBX5 | 26603904 |
| Cardiomyocyte | GATA4 | 26603904 |
| Cardiomyocyte | HAND2 | 26603904 |
| Cardiomyocyte | MIR590 | 27930352 |
| Cardiomyocyte | ZNF281 | 28982760 |
| Cardiomyocyte | MIR208B | 34426676 |
| Cardiomyocyte | SMAD6 | 37421991 |
| Cardiomyocyte | TBX20 | 37421991 |
| Hepatocyte | HNF1B | 23871605 |
| Hepatocyte | HNF6 | 24582926 |

| | | |
|---|---|---|
| Hepatocyte | PROX1 | 24582926 |
| Hepatocyte | MYC | 24582926 |
| Hepatocyte | ATF5 | 24582926 |
| Hepatocyte | HNF1A | 24582927 |
| Hepatocyte | FOXA1 | 24963715 |
| Hepatocyte | SOX2 | 26561000 |
| Hepatocyte | NR1I2 | 26561000 |
| Hepatocyte | HNF4A | 26603904 |
| Hepatocyte | FOXA3 | 26603904 |
| Hepatocyte | CEBPA | 26603904 |
| Hepatocyte | GATA4 | 26603904 |
| Hepatocyte | KDM2B | 27103435 |
| Hepatocyte | GATA3 | 28587331 |
| Hepatocyte | OCT4 | 28587331 |
| Hepatocyte | FOXA2 | 29192290 |
| Hepatocyte | KLF4 | 30635054 |
| Pancreatic cell | MIR302 | 25268319 |
| Pancreatic cell | MAFA | 25397325 |
| Pancreatic cell | NEUROD1 | 25397325 |
| Pancreatic cell | MAFB | 26603904 |
| Pancreatic cell | PDX1 | 26603904 |
| Pancreatic cell | PAX4 | 26603904 |
| Pancreatic cell | NEUROG3 | 26603904 |
| Pancreatic cell | PAX6 | 27833043 |
| Pancreatic cell | TGIF2 | 28193997 |
| Pancreatic cell | NKX2-2 | 36964318 |
| Neuron | OLIG2 | 20107439 |
| Neuron | DLX2 | 20502524 |
| Neuron | NEUROD1 | 21617644 |
| Neuron | LMX1A | 21725324 |
| Neuron | NEUROD2 | 21753754 |
| Neuron | MYT1L | 21802386 |
| Neuron | ISL1 | 21852222 |
| Neuron | EN1 | 22019014 |
| Neuron | ID1 | 22064700 |
| Neuron | NURR1 | 22105488 |

| | | |
|---|---|---|
| Neuron | PITX3 | 22105488 |
| Neuron | FOXA2 | 22426197 |
| Neuron | LMX1B | 23530235 |
| Neuron | OTX2 | 23530235 |
| Neuron | EMX2 | 24128663 |
| Neuron | POU3F4 | 24887289 |
| Neuron | SOX10 | 25158936 |
| Neuron | BCL11B | 25374357 |
| Neuron | DLX1 | 25374357 |
| Neuron | MIR9 | 25374357 |
| Neuron | MIR124 | 25374357 |
| Neuron | KLF7 | 25420066 |
| Neuron | NEUROG1 | 25420066 |
| Neuron | HES3 | 25454632 |
| Neuron | NOTCH1 | 25454632 |
| Neuron | DLX5 | 26526726 |
| Neuron | LHX6 | 26526726 |
| Neuron | LHX2 | 26603904 |
| Neuron | TCF3 | 26603904 |
| Neuron | PAX6 | 26603904 |
| Neuron | FOXG1 | 26603904 |
| Neuron | SOX2 | 26603904 |
| Neuron | ZIC1 | 26603904 |
| Neuron | REST | 26603904 |
| Neuron | POU3F2 | 26603904 |
| Neuron | HES1 | 26603904 |
| Neuron | RFX4 | 26603904 |
| Neuron | NEUROG2 | 26603904 |
| Neuron | ASCL1 | 26603904 |
| Neuron | PLAGL1 | 26603904 |
| Neuron | MYC | 26603904 |
| Neuron | KLF4 | 26603904 |
| Neuron | ISL2 | 26603904 |
| Neuron | LHX3 | 26603904 |
| Neuron | SOX11 | 26725112 |
| Neuron | HB9 | 28099929 |

| | | |
|---|---|---|
| Neuron | MIR218 | 28398344 |
| Neuron | GATA3 | 28587331 |
| Neuron | HMGA2 | 28844127 |
| Neuron | LHX1 | 28886366 |
| Neuron | MIR34B | 29526736 |
| Neuron | MIR34C | 29526736 |
| Neuron | PTF1A | 30030434 |
| Neuron | MSI1 | 31196173 |
| Neuron | MBD2 | 31196173 |
| Neuron | PHOX2B | 31315047 |
| Neuron | TFAP2A | 31315047 |
| Neuron | HAND2 | 31315047 |
| Paneth cell | HNF4A | 28943029 |
| Paneth cell | FOXA3 | 28943029 |
| Paneth cell | GATA6 | 28943029 |
| Paneth cell | CDX2 | 28943029 |
| Paneth cell | FOXA1 | 28943029 |
| Paneth cell | FOXA2 | 28943029 |
| Paneth cell | GATA4 | 28943029 |
| Skeletal muscle | MYOD1 | 3690668 |
| Skeletal muscle | MYCL | 28501623 |
| Skeletal muscle | MEF2B | 28808339 |
| Skeletal muscle | PITX1 | 28808339 |
| Skeletal muscle | PAX3 | 28808339 |
| Skeletal muscle | PAX7 | 28808339 |

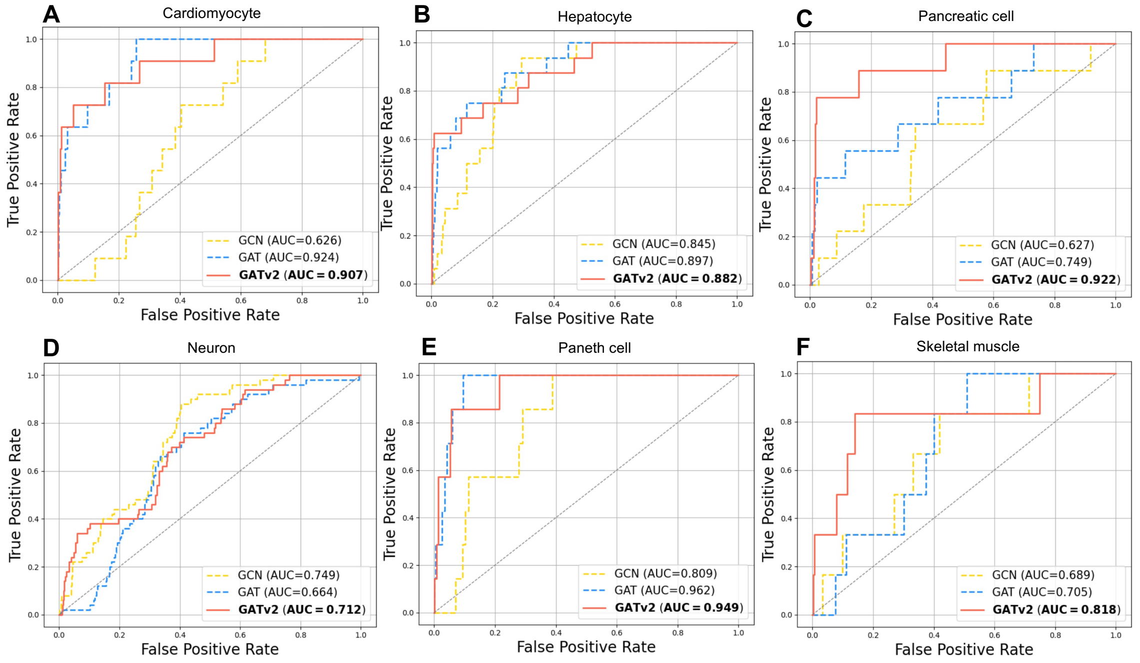


**Supplementary Figure 1.** Comparison of prediction performance across GNN architectures. (A–F) Comparison of prediction performance among GCN, GAT, and GATv2. ROC curves show the accuracy of predicting DR-inducing TFs during conversion from fibroblasts to (A) cardiomyocytes, (B) hepatocytes, (C) pancreatic cells, (D) neurons, (E) Paneth cells, and (F) skeletal muscle cells. Yellow and blue dashed lines indicate GCN and GAT, respectively, whereas orange solid lines indicate GATv2 (proposed method).

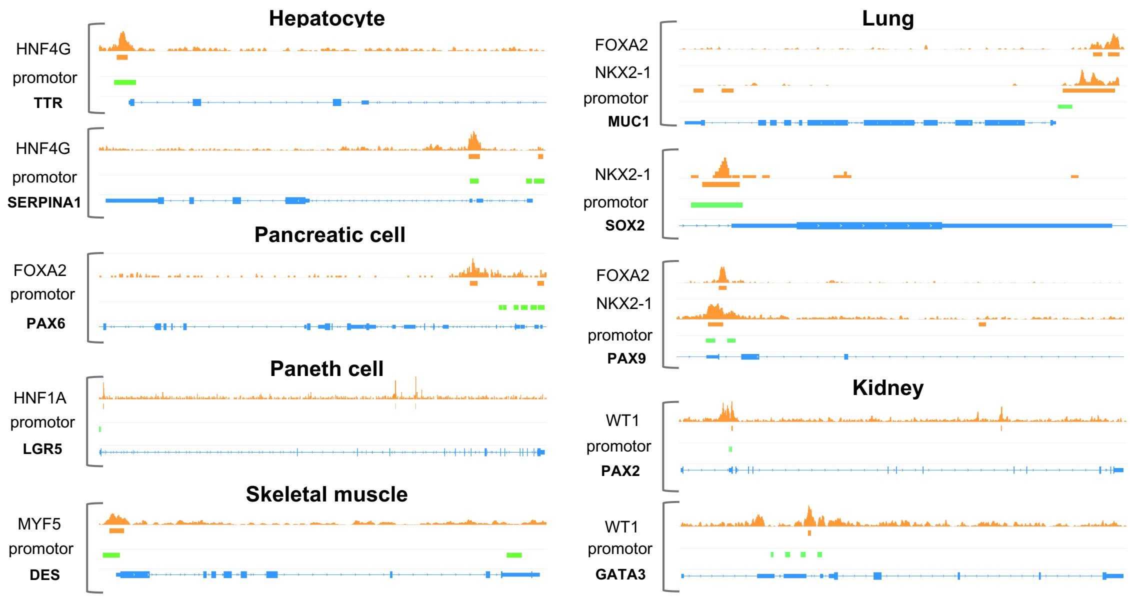


**Supplementary Figure 2.** Binding of predicted TFs to cell type–specific genes. ChIP-seq peaks of predicted TFs near promoter regions of cell type–specific genes. Orange peaks indicate TF binding signals, blue regions represent gene structures, and green regions denote promoter regions obtained from the ENCODE database [1].

**Supplementary Table 2.** Predicted DR-inducing TFs for tissues lacking reported DR-inducing factors. Top 20 TFs predicted by the proposed method for 11 tissues without previously reported DR-inducing TFs, ranked by prediction score.

| Rank | Kidney | Score | Lung | Score | Thyroid | Score | Adrenal gland | Score | Cervix | Score | Colon | Score |
|---|---|---|---|---|---|---|---|---|---|---|---|---|
| 1 | LHX1 | 0.944 | TBX4 | 0.873 | FOXE1 | 0.781 | NR0B1 | 0.942 | EVX2 | 0.474 | CDX1 | 0.908 |
| 2 | PAX2 | 0.939 | FOXJ1 | 0.704 | NKX2-3 | 0.654 | NR1H4 | 0.911 | EVX1 | 0.458 | CDX2 | 0.894 |
| 3 | UNCX | 0.934 | NKX2-1 | 0.693 | PAX8 | 0.618 | NR5A1 | 0.898 | HOXD12 | 0.427 | TBX10 | 0.876 |
| 4 | HNF1A | 0.930 | FOXF1 | 0.627 | NKX2-1 | 0.541 | CREB3L3 | 0.836 | HOXD13 | 0.418 | NKX2-3 | 0.866 |
| 5 | NR1H4 | 0.929 | FOXA2 | 0.620 | LHX8 | 0.529 | ZFP92 | 0.830 | HOXD11 | 0.413 | NR1I2 | 0.861 |
| 6 | HNF1B | 0.927 | CEBPE | 0.597 | TBX22 | 0.497 | PHOX2A | 0.772 | HOXA11 | 0.369 | NR1H4 | 0.859 |
| 7 | POU3F3 | 0.923 | TFEC | 0.559 | NKX2-8 | 0.416 | NR0B2 | 0.749 | HOXA13 | 0.362 | ATOH1 | 0.819 |
| 8 | WT1 | 0.916 | SPIC | 0.559 | GLIS3 | 0.387 | FEV | 0.734 | HOXD10 | 0.338 | SPIB | 0.818 |
| 9 | DMRT2 | 0.913 | LHX9 | 0.551 | NKX6-1 | 0.376 | SOX3 | 0.723 | LHX1 | 0.323 | CREB3L3 | 0.804 |
| 10 | SIM2 | 0.910 | ZNF705A | 0.538 | GCM2 | 0.375 | PHOX2B | 0.704 | HOXD9 | 0.318 | HNF1A | 0.800 |
| 11 | PAX8 | 0.910 | IRF8 | 0.486 | TBXT | 0.293 | GATA6 | 0.676 | PAX2 | 0.306 | HNF4A | 0.796 |
| 12 | HNF4A | 0.909 | SPIB | 0.482 | FOXA2 | 0.283 | EVX1 | 0.627 | MEIS3 | 0.251 | HNF4G | 0.795 |
| 13 | HNF4G | 0.908 | EOMES | 0.480 | DMRTA1 | 0.266 | LHX9 | 0.617 | NKX3-2 | 0.248 | HOXD12 | 0.757 |
| 14 | CREB3L3 | 0.890 | STAT4 | 0.479 | POU1F1 | 0.247 | RFX6 | 0.610 | FOXB2 | 0.247 | FOXF1 | 0.742 |
| 15 | HOXD11 | 0.886 | IRF7 | 0.458 | ARX | 0.247 | MYT1 | 0.600 | HOXD3 | 0.243 | NEUROG3 | 0.734 |
| 16 | FOXI1 | 0.878 | IRF5 | 0.446 | NKX2-4 | 0.246 | GATA4 | 0.597 | HOXA10 | 0.243 | HOXD13 | 0.732 |
| 17 | HMX2 | 0.868 | BATF2 | 0.444 | CUX2 | 0.244 | PEG3 | 0.596 | NKX2-6 | 0.241 | HOXB9 | 0.731 |
| 18 | NR0B2 | 0.867 | IRF1 | 0.421 | PAX2 | 0.241 | NEUROD4 | 0.593 | HOXD4 | 0.239 | PAX4 | 0.715 |
| 19 | EMX1 | 0.852 | KLF2 | 0.412 | SOX3 | 0.229 | HMX3 | 0.540 | NR0B1 | 0.234 | FEV | 0.704 |
| 20 | HOXB9 | 0.816 | GFI1 | 0.407 | HHEX | 0.228 | NEUROG3 | 0.528 | EMX2 | 0.234 | FOXA2 | 0.700 |

| Rank | Esophagus | Score | Ovary | Score | Prostate | Score | Spleen | Score | Stomach | Score |
|---|---|---|---|---|---|---|---|---|---|---|
| 1 | LHX8 | 0.345 | ZFP92 | 0.762 | EVX2 | 0.871 | SPIC | 0.832 | BHLHA15 | 0.952 |
| 2 | PAX9 | 0.283 | LHX9 | 0.677 | HOXD13 | 0.814 | SPIB | 0.824 | GATA4 | 0.932 |
| 3 | NKX2-3 | 0.255 | NR0B1 | 0.640 | EVX1 | 0.803 | CEBPE | 0.804 | RFX6 | 0.908 |
| 4 | NKX2-8 | 0.230 | LHX8 | 0.605 | NKX3-1 | 0.761 | NKX2-3 | 0.802 | FOXA2 | 0.891 |

| | | | | | | | | | | |
|---|---|---|---|---|---|---|---|---|---|---|
| 5 | NKX6-1 | 0.219 | NR1H4 | 0.546 | HOXD12 | 0.709 | EOMES | 0.783 | BARX1 | 0.889 |
| 6 | NKX2-6 | 0.214 | ARX | 0.534 | SOX14 | 0.662 | TLX1 | 0.775 | PDX1 | 0.880 |
| 7 | DMRTA2 | 0.210 | PEG3 | 0.528 | HOXA13 | 0.652 | IRF8 | 0.775 | RBPJL | 0.844 |
| 8 | PHOX2A | 0.197 | ZFPM2 | 0.473 | HOXD11 | 0.620 | TFEC | 0.763 | FOXA3 | 0.840 |
| 9 | SIM2 | 0.193 | GATA4 | 0.471 | HMX2 | 0.576 | PAX5 | 0.762 | PHOX2B | 0.809 |
| 10 | NKX3-2 | 0.191 | FOXL2 | 0.451 | LHX8 | 0.500 | NKX2-5 | 0.761 | HNF1B | 0.800 |
| 11 | TBX20 | 0.187 | NR5A1 | 0.445 | HOXD10 | 0.497 | IKZF3 | 0.752 | PTF1A | 0.788 |
| 12 | FOXF1 | 0.167 | DMRTC1B | 0.414 | SIM2 | 0.490 | STAT4 | 0.751 | HNF1A | 0.788 |
| 13 | PHOX2B | 0.160 | LHX1 | 0.396 | TBX4 | 0.478 | WT1 | 0.749 | HNF4A | 0.782 |
| 14 | FOXG1 | 0.153 | FIGLA | 0.388 | FEV | 0.451 | GFI1 | 0.746 | NKX2-3 | 0.769 |
| 15 | FOXF2 | 0.150 | ZNF275 | 0.383 | HOXD9 | 0.446 | TBX21 | 0.745 | HNF4G | 0.767 |
| 16 | CSRP3 | 0.147 | GATA6 | 0.379 | HOXA11 | 0.438 | ZNF705A | 0.743 | FEV | 0.754 |
| 17 | NKX6-2 | 0.146 | PAX2 | 0.354 | FOXB2 | 0.429 | LYL1 | 0.740 | NKX6-2 | 0.720 |
| 18 | TBX5 | 0.146 | SOX3 | 0.339 | PAX2 | 0.425 | BHLHA15 | 0.733 | ARX | 0.702 |
| 19 | FOXB2 | 0.140 | HOXD3 | 0.327 | CDX2 | 0.422 | TBXT | 0.732 | GATA5 | 0.689 |
| 20 | MYF6 | 0.140 | ZIM2 | 0.318 | BHLHA15 | 0.418 | IRF5 | 0.729 | PHOX2A | 0.665 |

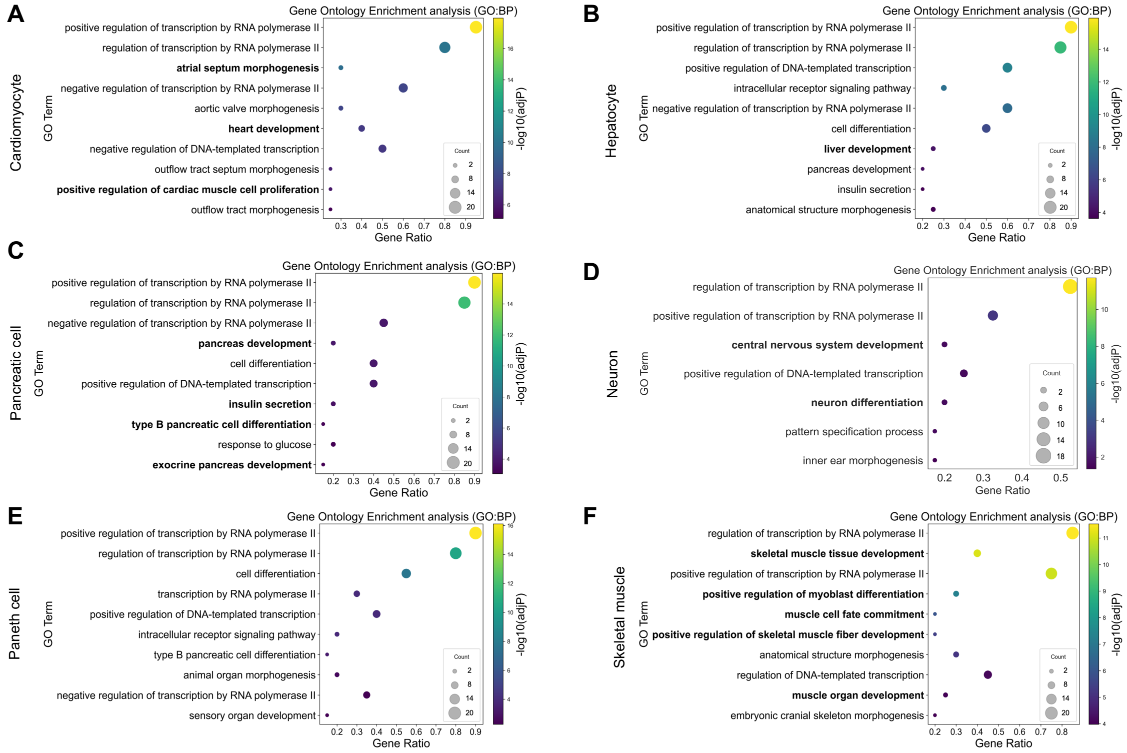


**Supplementary Figure 3.** Functional evaluation of predicted TFs via GO enrichment analysis. (A–F) Bubble plots showing GO enrichment analysis of the top 20 TFs predicted to induce DR from fibroblasts to (A) cardiomyocytes, (B) hepatocytes, (C) pancreatic cells, (D) neurons, (E) Paneth cells, and (F) skeletal muscle cells. Bubble color represents $-\log_{10}(adjP)$, bubble size indicates the number of input genes associated with each GO term, and the x-axis shows the proportion of input genes associated with each term. GO terms shown in bold are linked to tissue-specific development.

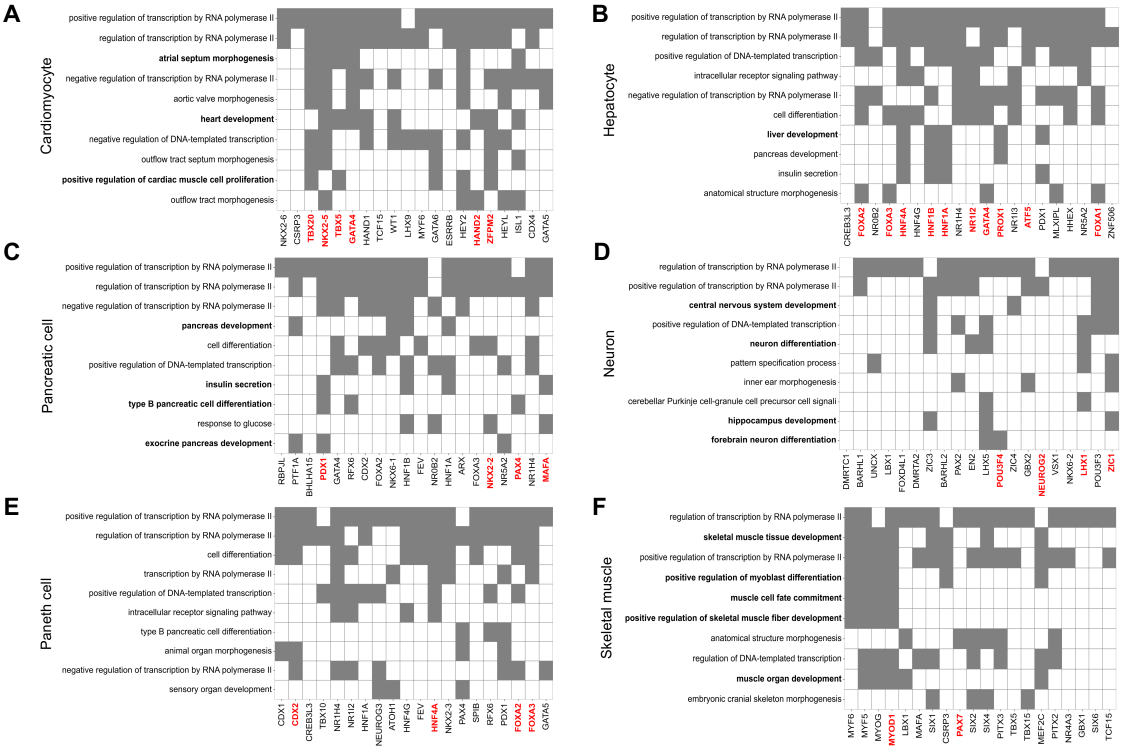


**Supplementary Figure 4.** Association between top-ranked predicted TFs and enriched GO terms. Correspondence between the top 20 TFs predicted to induce DR from fibroblasts to (A) cardiomyocytes, (B) hepatocytes, (C) pancreatic cells, (D) neurons, (E) Paneth cells, and (F) skeletal muscle cells and enriched GO terms. The x-axis lists the top-ranked TFs. Gray tiles indicate inclusion of a TF in the corresponding GO term. GO terms in bold are related to tissue-specific development, and those shown in red indicate known DR-inducing TFs.

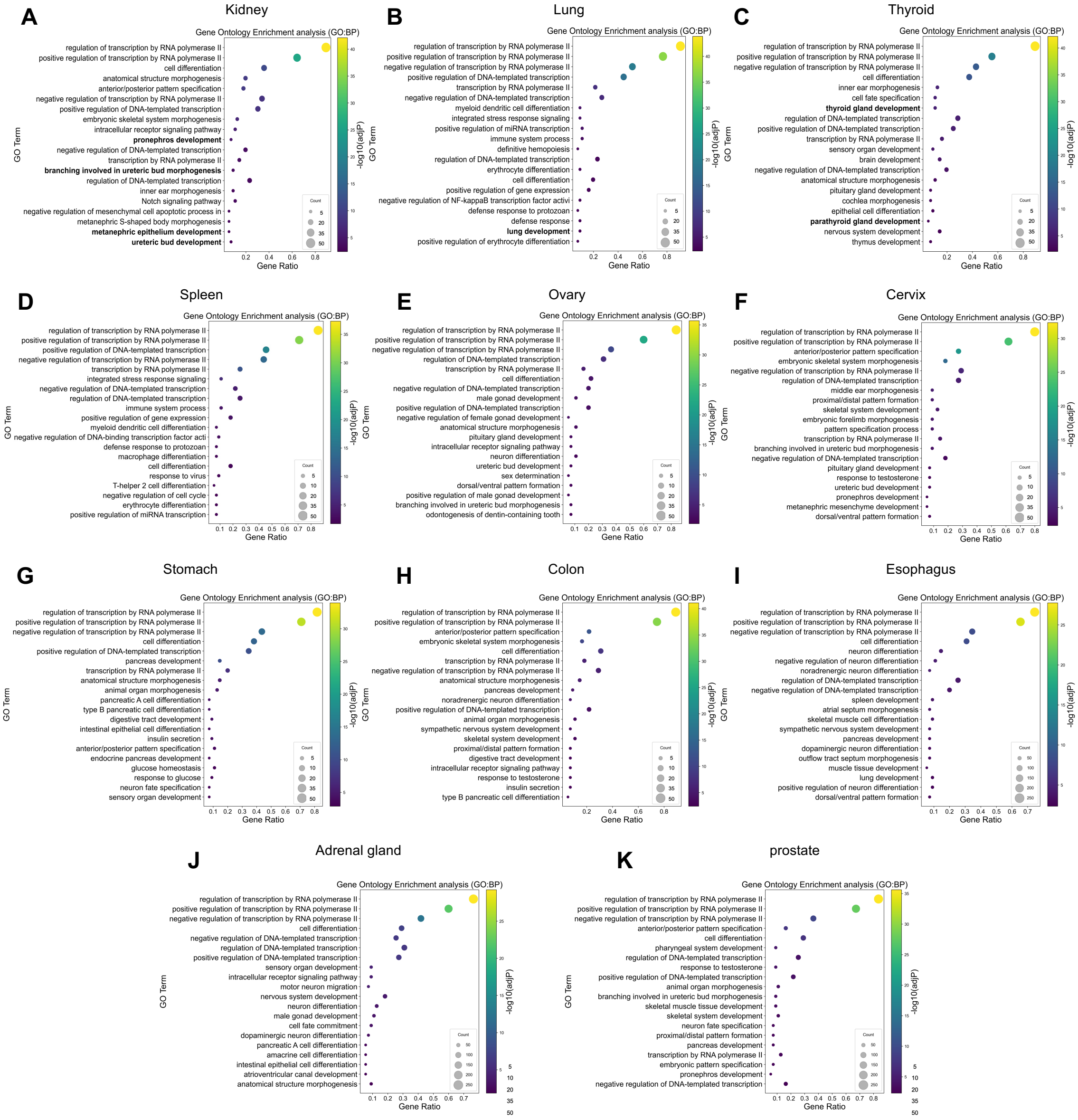


**Supplementary Figure 5.** Functional evaluation of predictions for tissues without reported DR-inducing TFs. Bubble plots of GO enrichment analysis for the top 5% of TFs predicted to induce DR from fibroblasts to (A) kidney, (B) lung, (C) thyroid, (D) spleen, (E) ovary, (F) cervix, (G) stomach, (H) colon, (I) esophagus, (J) adrenal gland, and (K) prostate tissues. Bubble color indicates $-\log_{10}(adjP)$, bubble size reflects the number of input genes per GO term, and the x-axis shows the proportion of input genes in each GO term. GO terms in bold are associated with tissue-specific development.

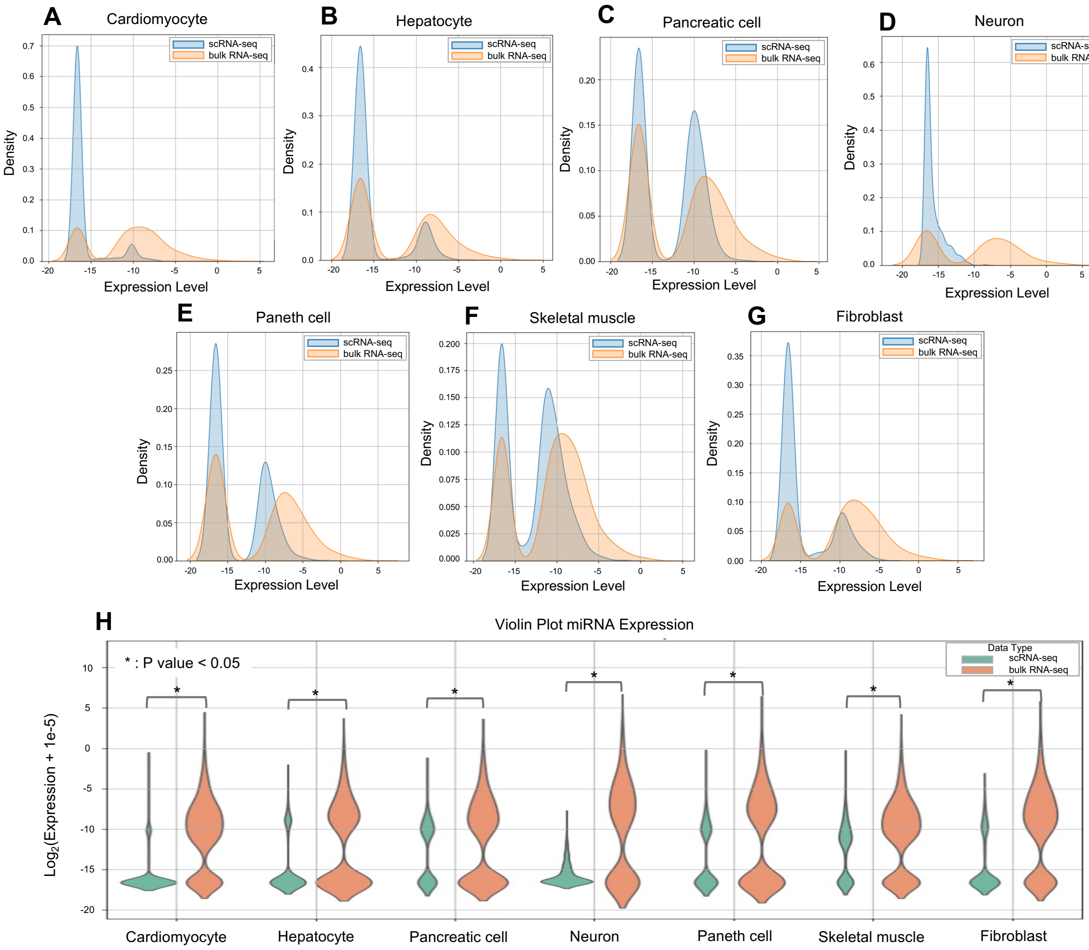


**Supplementary Figure 6.** Comparison of miRNA expression levels between single-cell and bulk RNA-seq. (A–G) Kernel density estimation (KDE) plots showing distributions of miRNA expression levels in (A) cardiomyocytes, (B) hepatocytes, (C) pancreatic cells, (D) neurons, (E) Paneth cells, (F) skeletal muscle cells, and (G) fibroblasts. The y-axis indicates probability density, and the x-axis shows miRNA expression levels. (H) Violin plots comparing miRNA expression levels across cell types. The y-axis indicates log-transformed values after adding 1e-5 to miRNA expression values. *: statistical significance at $p < 0.05$ (one- tailed *t*-test).

**Supplementary Table 3.** Predicted DR-inducing factors including miRNAs. Top 20 TFs and miRNAs predicted via the proposed method for the 6 cell types with reported DR-inducing TFs. Known DR-inducing TFs and miRNAs are shown in bold.

| Rank | Cardiomyocyte | Score | Hepatocyte | Score | Pancreatic cell | score | Neuron | Score | Paneth cell | Score | Skeletal muscle | Score |
|---|---|---|---|---|---|---|---|---|---|---|---|---|
| **1** | NKX2-6 | 0.916 | MIR4539 | 0.990 | RBPJL | 0.960 | MIR153-1 | 0.988 | CDX1 | 0.954 | MIR4269 | 0.956 |
| **2** | CSRP3 | 0.915 | MIR587 | 0.989 | PTF1A | 0.951 | MIR4675 | 0.923 | MIR509-2 | 0.951 | MYF6 | 0.956 |
| **3** | **TBX20** | 0.915 | MIR7155 | 0.988 | BHLHA15 | 0.949 | MIR4674 | 0.915 | MIR548H3 | 0.951 | MYF5 | 0.954 |
| **4** | **NKX2-5** | 0.912 | MIR3683 | 0.987 | **PDX1** | 0.942 | MIR8088 | 0.911 | MIR548I3 | 0.938 | MYOG | 0.954 |
| **5** | **TBX5** | 0.909 | CREB3L3 | 0.986 | GATA4 | 0.937 | MIR378B | 0.908 | MIR4321 | 0.937 | **MYOD1** | 0.952 |
| **6** | MIR4460 | 0.909 | MIR670 | 0.986 | MIR6507 | 0.935 | MIR384 | 0.903 | MIR509-1 | 0.933 | LBX1 | 0.951 |
| **7** | **GATA4** | 0.906 | MIR216B | 0.986 | ONECUT1 | 0.932 | MIR5682 | 0.902 | **CDX2** | 0.933 | MAFA | 0.951 |
| **8** | MIR5589 | 0.904 | **FOXA2** | 0.986 | RFX6 | 0.932 | MIR572 | 0.900 | CREB3L3 | 0.931 | SIX1 | 0.951 |
| **9** | MIR5689 | 0.902 | NR0B2 | 0.986 | CDX2 | 0.925 | DMRTC1 | 0.900 | MIR5692A2 | 0.926 | MIR3163 | 0.951 |
| **10** | HAND1 | 0.901 | **FOXA3** | 0.985 | FOXA2 | 0.924 | BARHL1 | 0.898 | MIR181A1 | 0.920 | MIR5689 | 0.950 |
| **11** | TCF15 | 0.899 | MIR502 | 0.985 | NKX6-1 | 0.916 | UNCX | 0.898 | TBX10 | 0.918 | MIR519E | 0.950 |
| **12** | MIR3163 | 0.896 | **ONECUT1** | 0.985 | HNF1B | 0.913 | LBX1 | 0.897 | MIR6724-4 | 0.918 | CSRP3 | 0.948 |
| **13** | MIR302E | 0.891 | **HNF4A** | 0.984 | FEV | 0.912 | FOXD4L1 | 0.897 | NR1H4 | 0.916 | MIR4692 | 0.948 |
| **14** | MIR8068 | 0.889 | MIR4478 | 0.984 | NR0B2 | 0.910 | DMRTA2 | 0.895 | NR1I2 | 0.914 | **PAX7** | 0.946 |
| **15** | MIR4727 | 0.887 | MIR8054 | 0.984 | HNF1A | 0.910 | ZIC3 | 0.894 | MIR551A | 0.913 | MIR7153 | 0.946 |
| **16** | MIR4692 | 0.871 | HNF4G | 0.983 | MIR217 | 0.908 | BARHL2 | 0.894 | HNF1A | 0.902 | MIR4727 | 0.945 |
| **17** | WT1 | 0.869 | MIR508 | 0.983 | ARX | 0.906 | PAX2 | 0.894 | NEUROG3 | 0.896 | MIR6086 | 0.943 |
| **18** | MIR633 | 0.862 | MIR618 | 0.983 | FOXA3 | 0.904 | EN2 | 0.892 | ATOH1 | 0.895 | MIR133B | 0.940 |
| **19** | MYOCD | 0.862 | **HNF1B** | 0.983 | **NKX2-2** | 0.903 | LHX5 | 0.892 | MIR520F | 0.895 | MIR548AP | 0.938 |
| **20** | LHX9 | 0.859 | **HNF1A** | 0.983 | NR5A2 | 0.901 | **POU3F4** | 0.892 | MIR758 | 0.892 | SIX2 | 0.937 |
